# Incorporating gene expression and environment for genomic prediction in wheat

**DOI:** 10.1101/2024.09.26.612369

**Authors:** Jia Liu, Andrew Gock, Kerrie Ramm, Sandra Stops, Tanya Phongkham, Adam Norman, Russell Eastwood, Eric Stone, Shannon Dillon

## Abstract

The adoption of novel molecular strategies such as genomic selection (GS) in crop breeding have been key to maintaining rates of genetic gain through increased efficiency and shortening the cycle of evaluation relative to conventional selection. In the search for improved methodologies that incorporate novel sources of variation for the assessment of genetic merit, GS remains a focus of crop breeding research globally. Here we explored the role transcriptome data could play in enhancing GS using wheat as a test case. Across 286 wheat lines, we integrated phenotype and multi-omic data from controlled environment and field experiments including ca. 40K single nucleotide polymorphisms (SNP), abundance data for ca. 50K transcripts as well as meta-data (e.g. categorical environments) predicted individual genetic merit for two agronomic traits, flowering time and height. We combined phenotype and multi-omic data from both controlled environments and field experiments. This included ca. 40K single nucleotide polymorphisms (SNPs), ca. 50K transcript abundance data, and metadata (such as categorical environmental conditions). Using this integrated data, we predicted individual genetic merit for two agronomic traits: flowering time and height. We evaluated the performance of different model scenarios based on linear (GBLUP) and Gaussian/nonlinear (RKHS) regression in the Bayesian analytical framework. These models explored the relative contributions of different combinations of explanatory variables; additive genomic (G), transcriptomic (T) and environment (E), with and without considering non-additive epistasis and the *G*×*E* random effects. In controlled environments, where traits were measured under contrasting daylength regimes (long and short days), transcriptome abundance outperformed other explanatory variables when considered independently, while the model combining SNP, environment and *G* × *E* marginally outperformed the transcriptome. The best performing model for prediction of both flowering and height combined all data types, *G* × *E* and epistasis, where the GBLUP framework showed slightly better performance overall compared with RKHS across all tests. Under field conditions, we similarly found that models combining all variables were superior, with the GBLUP and RKSH methods performing equally well. However, the relative contribution of the transcriptome was reduced. Our results show there is a predictive advantage to direct inclusion of the transcriptome for genomic evaluation in wheat breeding. However, the complexity and cost of generating large scale transcriptome data are likely to limit its feasibility for commercial breeding. We demonstrate that combining less costly environmental covariates with conventional genomic data provide a practical alternative with similar gains to the transcriptome when environments are well characterised.

**Highlights:**

- Incorporating transcriptome and environment in genomic prediction;
- Model comparisons and Bayesian inference.
- Differential random effects of transcriptome, SNP and environment.

## 1 Introduction

Over the last decade genomic selection (GS, Meuwissen et al. (2001)) has provided significant advances in animal and plant breeding, by allowing breeders to efficiently identify and select individuals with the highest genetic merit in relation to a trait of interest. These approaches work best for highly heritable traits that are complex in their genetic control. GS leverages genomic information to predict the breeding value (GEBVs) of individuals in a population, which are usually predict from high-density single nucleotide polymorphism (SNP) markers and observed training data via a statistical model.

This model fits the relationship between genotypes (SNP) and phenotypic trait(s) based on training population data and partitions the contribution of different genomic effects (e.g. genomic additive and non-additive effects such as epistasis and dominance) on trait variance. Traditional statistical approaches for GS include genomic best linear unbiased prediction (GBLUP, Clark and van der Werf (2013)) and Reproducing kernel Hilbert space regression (RKHS, see e.g., Gianola and Van Kaam (2008)), a non-linear Gaussian kernel regression model. GBLUP is based on the linear mixed model, it typically uses genomic SNP markers to capture genomic effects from well-defined fixed and random components (Mardia et al., 2024) and could be considered a gold-standard for GS. Because of its non-linearity, RKHK can be used to effectively capture a combination of complex genomic effects from low- and high-order perspectives, see e.g., Heffner et al. (2009).

The potential for other omics data types (e.g. transcriptome, proteome, metabolome) to improve the accuracy of GS in crops has recently gained attention (Li et al., 2019). These biological data layers sit functionally between the genome and phenotype expression and can be considered a molecular proxy for phenotype, or endophenotype (Te Pas et al., 2017). Some studies have shown the value of integrating omic data through prior analysis to resolve the biological mechanisms driving variation in phenotype, and based on this inform which parts of the genome receive greater attention in the GS model, e.g. by weighting gene based markers (Fang et al., 2017; Ye et al., 2020). This approach is underpinned by detailed experimental work and an in-depth understanding of the trait biology and genetics to guide model development, with a distinct risk that biases in interpretation will be propagated into predictions.

A data-driven alternative is to include multi-omic data directly in the predictive framework. Typically GS is conducted with sparse genomic SNP data that exploits linkage to capture global genomic variation. Alternate data layers such as the transcriptome may provide one avenue to improve the density of biologically relevant markers by focusing on functionally active regions of the genome. It has also been proposed that omics data layers such as the transcriptome could improve GS predictions by more effectively capturing computationally elusive epistatic interactions. Computing higher order interactions among 10’s of thousands to millions of SNP markers rapidly becomes intractable. Whereas the transcriptome provides a biologically informed dimensionality reduction since epistatic interactions among multiple genomic loci could have an additive effect on transcript abundance (Li et al., 2019). In a mechanistic sense the transcriptome sits functionally between the genome and phenotype, indirectly capturing both genetics (G) and environmental (E) effects and their interactions (*G* × *E*). Where *G* × *E* plays an important role in mediating trait expression, inclusion of transcriptome information provides one avenue to capture this in prediction frameworks, in particular where environmental effects are not well characterised.

The majority of studies investigating the utility of transcriptomes in GS have been in maize (Frisch et al., 2009; Fu et al., 2011; Guo et al., 2016; Zenke-Philippi et al., 2017; Xu et al., 2017; Schrag et al., 2018; Westhues et al., 2017; Wang et al., 2019; Azodi et al., 2019). Largely, these illustrate that the transcriptome provides equal or better prediction accuracy than the genome alone. By combining additional omic strata, prediction accuracy can often be improved further, with performance varying slightly depending on the choice of prediction algorithm. Though there are exceptions: For example, Xu et al. (2017) found that genome SNP data was a better predictor of yield traits (e.g. ear length, weight) than transcriptome or metabolome data layers in maize. This result potentially points to opportunistic use of omic data where the tissue and time point for sample collection (immature seed) were not optimised for predicting yield traits in the field at maturity. This points to a significant challenge in implementing large scale transcriptome studies for trait prediction, where careful factoring of temporal, developmental and environmental cues into sampling of endophenotypes for trait prediction is needed. Further, in applied settings it will be desirable to integrate ‘omics data collected in the field for GS, where this challenge increases due to greater temporal environmental variability. Despite being a potentially important question to resolve, there are few studies to date that explored the use of endophenotypes under field conditions for GS applications, including feasibility in commercial breeding programs, suggesting more work is needed in a broader range of crops. While there are many studies in maize, there are not in wheat. Significantly, the application of multi-omics for GS in wheat, an important staple crop in Australia and globally, has until today not been explored.

Productivity of wheat has long been maximised by breeders and growers optimising the timing of flowering to match local climate (Reynolds et al., 2012; Hyles et al., 2020). This has been achieved directly and indirectly by selecting genetic and environmental drivers of flowering and their interactions (*G* × *E*), some of which are now well characterised (Crossa et al., 2021; Tolhurst et al., 2022). Flowering time in wheat could therefore be a suitable model for exploring the relative contributions of genomic, endophenomic and environmental effects and their interactions on trait variation. Further to this, as a deterministic trait the molecular control of flowering is decided early in development, and patterns of regulatory expression are detectable before floral transition, see VanGessel et al. (2022); Shi et al. (2019). Early selection is desirable in breeding and flowering could provide a useful platform to explore whether endophenotypes sampled at early stages of development can robustly predict traits which manifest later in plant development. Environments experienced at all growth stages can shift variation in phenology via *G* × *E*, thus another important question is the extent to which such variation impacts the efficacy of endophenotypes for GS in field relative to static conditions in controlled environments. This will be particularly relevant for broad acre, dry-land crops such as wheat. Lastly, given endophenotypes are an expression of underlying genetics and environment, something that has received less attention in the literature is whether they can more efficiently be represented by robustly capturing G, Epistasis and *G* × *E* interactions in prediction frameworks.

This study evaluates two widely used genomic selection approaches to predict agronomic traits in wheat using a diverse wheat panel and data collected in both field and controlled environments. Specifically, we use a linear mixed model, GBLUP, as a benchmark to compare with a non-linear Gaussian kernel regression (RKHS) under the Bayesian framework, and compare different model scenarios designed to test the relative merits of applying different combinations of predictor variables; additive genomic (G), transcriptomic (T) and environmental (E) covariates, with and without considering non-additive random genomic effects (epistasis and dominant) and *G* × *E*. We highlight the value of including transcriptome for prediction and its potential for application in wheat breeding. We also consider the challenges associated with field-based transcriptome wide experiments, including choice of endophenotype sampling in settings where environment is likely to vary significantly during plant development. Considering such challenges and greater cost of data generation we ask the question, “is this approach feasible for commercial breeding programs”, and explore the role of lower cost alternatives, such as including *G* × *E*, in supporting improvements in GS.

## 2. Data and Experiments

### 2.1. Genetic material

This study used the OzWheat diversity panel, a collection of 286 wheat lines that include land races and progenitors of early Australian varieties, additional founders that emerged through the green revolution, and a larger number of Australian modern elite varieties (Hyles et al., 2024; Dillon et al., 2024). In this study a subset of 286 lines from OzWheat Panel (Hyles et al., 2024) were used in controlled environment and field experiments. Across field experiments there were slight variations in which of the 286 lines were used depending on quantities of seed available at the time of sowing.

### 2.2. Controlled environment experiments

All data collected in controlled environments were as reported in detail by Dillon et al. (2024). In breif, panel genotypes were grown under contrasting ‘long’ (16hr light) and ‘short’ (8hr light) photoperiod in controlled-environment growth chambers (PGC20 Conviron®, Winnipeg, Canada). For each variety there were six biological replicates in the experiment. All of these were analysed. Panel genotypes were grown under contrasting ‘long’ (12hr light) and ‘short’ (8hr light) photoperiods in a doubled coated plastic growth house with temperature control in Canberra.

#### Trait data

Plants were subject to twice weekly assessments to detect flowering. This was based on plants having reached stage z61 or “anthesis” (days after sowing), marked by the extrusion of anthers from the spikelets. Height (cm) of each plant was measured at maturity, and included the total above ground stem length, plus total spike length. Both of these traits are highly heritable with strong genetic control, and in the case of flowering exhibit strong environmental and *G* × *E* interactions with photperiod variation.

#### Transcriptome data

The crown plus coleoptile was harvested at the 2-leaf stage (Z12) from all seedlings grown in the cabinet experiment, which were immediately stored in pre-labelled tubes and frozen in liquid nitrogen. Samples were subsequently transferred to -80°C for long term storage. Sample collection was timed to occur over the two hours up to midday in each treatment (long and short daylengths). Cabinet time of day was staggered by 2 hours between treatments to allow for both treatments to be sampled on the same day. Ribonucleic acid (RNA) was extracted from entire frozen tissue sampled from a single biological replicate of each panel variety and libraries prepared for RNA-seq. Sequenced reads were quality checked, trimmed and mapped against the Chinese Spring reference coding sequence v1.0 using the Trinity package (Haas et al., 2013), for estimation of expression abundance for 44054 coding genes.

#### Genome SNP data

Single nucleotide polymorphsim (SNP) data was obtained from two sources. Trimmed paired end sequence reads for each sample were merged across treatments and aligned to the Chinese Spring coding sequence (CDS) reference v1.0 (IWGSC et.al 2014) using BWA-MEM (Li (2013), settings) and SNP variants were called using GATK3.7 haplotype caller as described by Dillon et al. (2024), yielding ca. 12K SNP markers. These were combined with ca. 21K SNPs from the 90K Illumina Infinium SNP array (Wang et al., 2014) to make up a total set of 33174 SNP markers for downstream analysis as described in Hyles et al. (2024).

### 2.3. Field experiments

In total four field experiments were conducted at the CSIRO Ginninderra Experimental Station (GES) near Canberra in 2018 (35°11’59”S 149°04’48”E) and 2019 (35°10’58”S 149°03’30”E), and at the Australian Grain Technologies (AGT) breeding site, Kabinga near Wagga Wagga in 2018 (35°03’28”S 147°02’44”E) and 2019 (35°03’29”S 147°02’51”E), previously described by Hyles et al. (2024). Two replicates of each line were sown in a randomised complete block design at each site (n=260 at Wagga, n=280 in Canberra). Experiments were sown at GES on the 5th of June and 30th of April, and on the 25th of May and 28th of May at Kabinga in 2018 and 2019 respectively. In Canberra, each plot comprised eight rows with 18 cm spacing and length of five linear metres. At Wagga Wagga plots comprised two rows only, with total plot dimensions 0.75 × 2.5 linear metres. Environmental covariates were not used to characterise experiments in our analysis, rather we treated site as a categorical variable in our models. Nevertheless, environmental conditions do varied significantly between our chosen locations. Canberra was consistently cooler through the growing season compared to Wagga Wagga over the years our experiments were conducted, as shown in Figure 1. The sites are similar in terms of latitude and hence photoperiod during the growing season.

**Figure 1.**
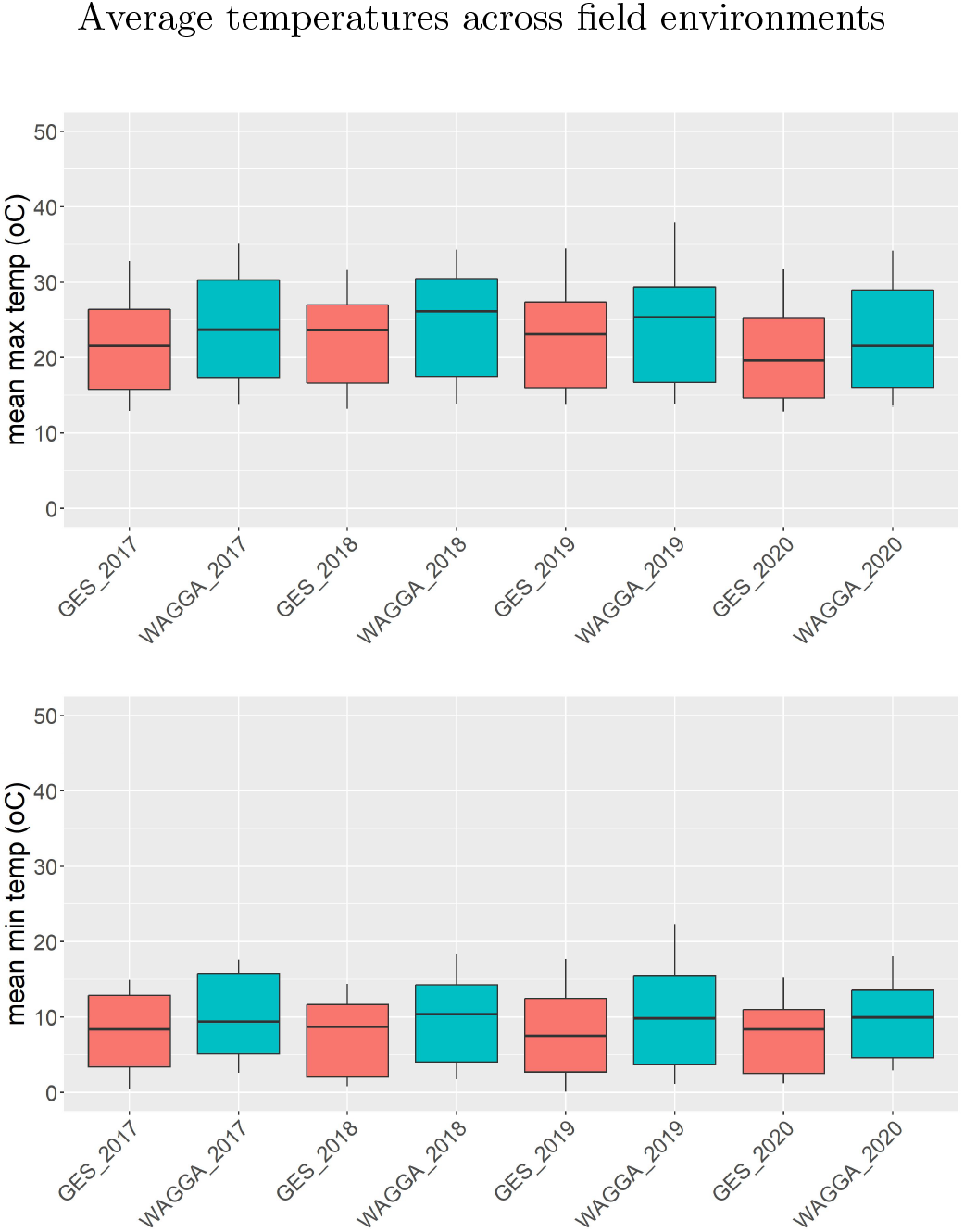
Means of maximum and minimum temperatures from five different field trials in Canberra and Wagga Wagga over four different years between 2017-2020.

#### Trait data

For each plot we obtained estimates of date to heading (Z51, date that 50% of plants in the plot had spikes fully emerged from the boot), with the exception of Wagga Wagga 2019, where heading date was only obtained on a single replicate block. Height (cm) at maturity measured for 3 representative plants per plot, and included the total above ground stem length, plus total spike length. Both of these traits are highly heritable with strong genetic control and in the case of flowering exhibit strong *G* × *E* interactions with thermal and vernal accumulation. The same technique of trait data collection as in Hyles et al. (2024).

#### Molecular data

The crown plus coleoptile was harvested at the 2-leaf stage (Z12) from 2 representative seedlings per field plot in the first block, which were immediately stored in pre-labelled tubes, frozen in liquid nitrogen and stored on dry ice for transport back to the laboratory. Samples were subsequently transferred to -80°C for long term storage. Sample collection was timed to occur over the two hours up to midday. RNA was extracted from frozen tissue using the Maxwell® RSC Plant automated extraction system following manufacturer’s instructions (Promega, catalogue number AS1500), and quality checked according to the method described above. RNA libraries were generated using the method of Wang et al, (2011) with modifications (supplemental file S1), as described in Dillon et al. (2024). The multiplexing design used 384 polymerase chain reaction (PCR) primer combinations to introduce the dual end 8-bp index sequence to the final library product using the TruSeq backbone, which was compatible with the Illumina Novaseq 6000 sequencing platform. Libraries from each experiment were sequenced on one lane of a Novaseq 6000 S4 flow cell. Using the same workflow as described above for the controlled environment experiments the abundance of 70,606 coding genes was obtained from the sequence data and represented as a sparse matrix for downstream analysis. Single nucleotide polymorphsim (SNP) data used in combination with the field trait data is the same as that described above for the controlled environment experiments.

## 3. Statistical models for genomic selection and prediction

### 3.1. GBLUP

The conventional genomic best linear unbiased prediction (GBLUP) model only considers the genomic additive random effect with a simple expression as the following,

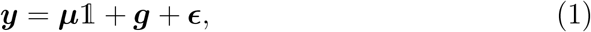

where ***y*** is the trait of interest, ***μ*** is *n* × 1 vector that describes the fixed effect, the noise term 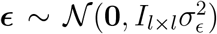. We have the genomic random 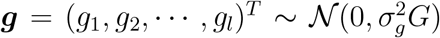 follows a multivariate Gaussian distribution (MVN) with zero mean and the covariance matrix as the product of from the genomic variance 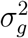 and the genomic relationship matrix (GRM), *G*, VanRaden (2008); Casella and Berger (2002), representing the covariance between pair-wide wheat lines (genotype). We use a method proposed by Endelman and Jannink (2012) to estimate GRM via the linear kernel 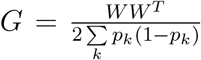, where *W* is the centered genotype matrix with *W*_*ik*_ = *X*_*ik*_ − 2*p*_*k*_, *k* = 1, 2, …, *m*, and *X* is a *l* × *m* matrix, containing genomic SNP markers with *l* individual lines and *m* bi-allelic (AA,AB and BB) SNP markers coded as [-1,0,1] or [0,1,2] at each line, *p*_*k*_ allele frequency equalling 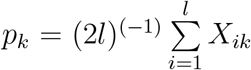 accounted from DNA SNP sequences for each individual line, see Müller et al. (2015). This setting ensures a proper scaling of the diagonal element of the estimated GAM equals 1 + *f*, where *f* is the inbreeding coefficient of the current population of interest. When *n* ≠ *l* the number of trait observations is not match the number of individual lines, an incidence matrix *Z*_*g*_ Li et al. (2019) (0 absence and 1 presence) may be introduced such that 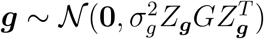 so that the genotypes relate to the phenotype observations. For examples, suppose *G* is a 3 × 3 GRM describes correlation between three individual lines, and we have 10 trait observations, 5 from the 1st line, 3 from the 2nd line and 2 from 3rd line, then we have an incidence matrix 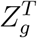 with the dimension 3 × 10 given by

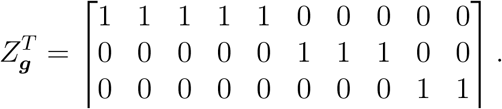

Non-additive genomic random effects including epistasis (or gene-by-gene, *G*#*G*) and dominant (A). Epistasis refers gene-gene interactions between loci, and can appear in biallelic and / or high order. Epsitasis has been reported to be able to modify phenotypic traits in crops and may have advantages for GS Doust et al. (2014). Whereas dominant effect describes interactions between alleles at the same locus, and typically refers heterozygous allels, AB. To analyse dominant effect the bi-allelic SNP markets, we can encode the SNP markers as 1 (AA, BB) and 0 (AB).

To integrate non-additive genomic random effects with the additive, the traditional GBLUP model in Equation 1 can be extended by

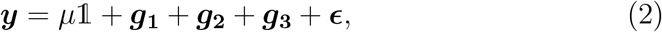

where ***g***_1_ is genetic additive random effect as ***g*** in Equation 1, ***g***_2_ stands for the epistasis and ***g***_3_ is the dominant effect. Due to the property of the epsitasis, we can assume ***g***_2_ follows a Gaussian distribution with zero means and a product of the linear kernel *G*, that is, 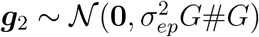, where # is the Kronecker product between GAM, *G*#*G* = *GG*^*T*^ − *diag*(*G*) and 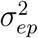 is the epistasis variance. The dominant random effect 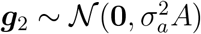 follows a MNV with zero means and covariance from dominant relation matrix *A* and the variance 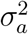. If *n* ≠ *l*, the same incidence matrices *Z*_*g*_ can be introduced to each random effect in Equation 2.

Li et al. (2019) have extended the GBLUP model to a GTBLUP model (3) by integrating omics transcriptome (T) data with a math model as below

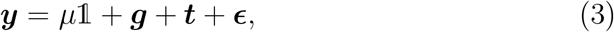

where the transcriptome effect indicates by ***t***, which can be assumed to be a MVN, 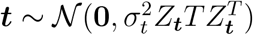 as ***g***.

The influence of gene-by-environment (G × E) in crops has been studied in recent decades(Doust et al., 2014; Jarquin et al., 2014; Bandeira e Sousa et al., 2017; Lopez-Cruz et al., 2015). G × E refers to certain situations where the relative allele effects vary across the environments. It has be claimed to have strong affect to some visible traits such as branching, seed size, etc (Sadras and Slafer, 2012). In this paper, we analyse two agronomic traits in wheat, height and flowering time, that are affected by multiple random effects introduced above. We hence extend the model in (3) by involving G × E,

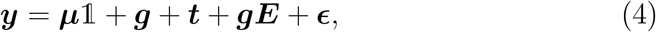

where ***g****E* represent interactions between genomic SNP markers and the environment, and 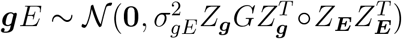, where *Z*_***E***_ is the incidence matrix for the effects of environments to the traits, and ∘ denote the Hadamard or Schur product, describing the element to element product between two matrices at same order Bandeira e Sousa et al. (2017); Jarquin et al. (2014), and 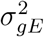 indicates the genomic over environmental variance.

### 3.2. RKHS

Reproducing kernel Hilbert space (RKHS) Berlinet and Thomas-Agnan (2004) introduces a kernel function: *k* : 𝒳 × 𝒳 → ℝ named a reproducing kernel over a non-empty feature set 𝒳 through a map *ϕ* : 𝒳 → ℋ over a Hilbert space ℋ such that

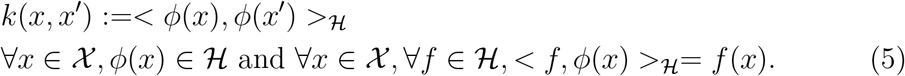

In the genomic prediction the RKHS model introduces a nonlinear Gaussian kerner on 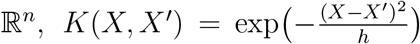 Li et al. (2019); Jarquin et al. (2014); Costa-Neto et al. (2021) to capture a mixed genomic random effects including additive as well as complex cryptic interactions, which we call it *epistasis*, between pair-wise SNP markers and also dominant effect (A). This is main different between the RKHS model and the GBLUP models, where in GBLUP a linear kernel is applied to capture additive (GAM) and non-additive (epsidasis and dominant) genomic random effects separately. We can also use this nonlinear kernel to describe other random effects in the model. *h* in the Gaussian kernel describes a bandwidth parameter controlling the decay rate of the correlations between individuals.

#### Modeling G × E

In the *j*th environment the model in Equation (4) can be extended by

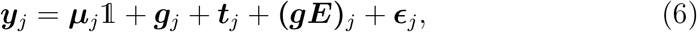

so that the observed trait data is reorganized based on *J* environments given by

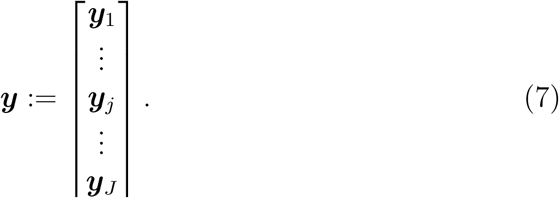

We apply the multi-environment, single variance model proposed by Bandeira e Sousa et al. (2017) to capture the G× E interactions by

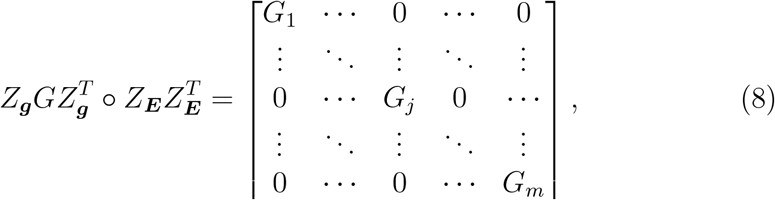

where *G*_*j*_ represents correlations between wheat lines in the *j*th environment. The main reason to apply this model is that our data were collected under environmental scenarios. Other possible G × E models can be found in e.g., Jarquin et al. (2014); Bandeira e Sousa et al. (2017); Lopez-Cruz et al. (2015). Assuming *X*_*j*_ is the input data at *j*t environment from an arbitrary random effect, the variance-corvariance matrix for ***g*** or ***t*** becomes

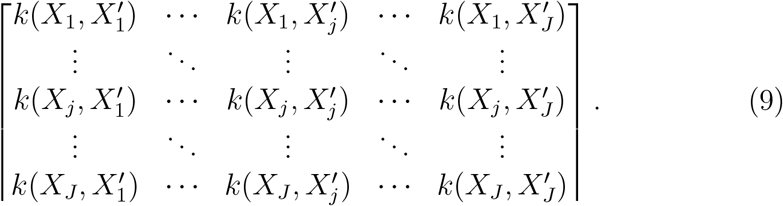

### 3.3. Bayesian inference for prediction

#### Bayesian theorem and the hierarchical model

Let **u** = be the unknown random effects of interest in the model, and the corresponding variance is denoted as *σ*_***u***_, and assuming ***ξ*** = {***μ***, *σ*_***u***_, ***θ***, *σ*_*ϵ*_} where ***θ*** includes all the possible hyperparameters in the kernel functions, our likelihood can be then expressed as

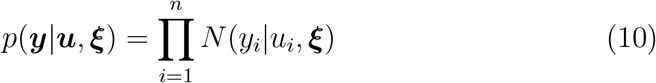

By Bayesian theorem the joint posterior can be approximated by

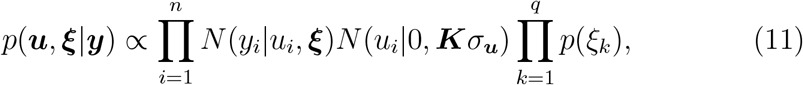

where *N* (*u*_*i*_|0, ***K****σ*_***u***_) is the Gaussian prior of the unknown random effects. The variance-covariance matrix ***K*** capture the correlations from different types of input data. The hyperprior density of the hyperparameter ***ξ*** can be expressed by

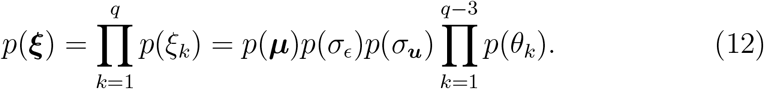

We optimise the model by maximizing the logarithmic posterior up o a constant w.r.t. ***ξ***,

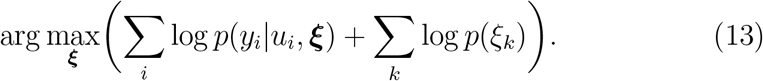

We then use the posterior predictive density which is a Gaussian from the trained model with optimal value 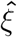 to do the prediction at new input data *X*^*^,

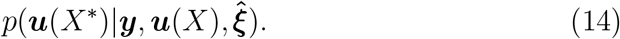

By choosing conjugate priors for the model parameters, e.g., *p*(***μ***) is constant, *p*(***u***) follows the multivariate Gaussian distributions, *p*(*σ*_*ϵ*_) and *p*(*σ*_***u***_) follow the scaled inverted *χ*^2^ distributions (see more details in Chapter 16 Mrode (2014)), we can have close forms of the full conditional posterior distributions of each parameter and hyperparameter of interest. The predictive values at new input data then can be computed from the posterior predictive distribution Equation (14), see for instance Rasmussen (2006). There are multiple ways to train the model, in this work we apply the Gibbs sampler though Markov chain Monte carlo method (MCMC) to optimize ***ξ***, see more details in e.g., Robert and Casella (2005).

#### The eigen-decomposition transformation

The eigen-decomposition is widely used in the computation for stability and efficiency by making sure the variance-covariance matrix is well-conditioned and symmetric. By the eigen-decomposition, we have

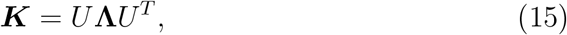

where *U* is a *n* × *n* square matrix whose *i*th column is the corresponding eigenvector of ***K***, and *U* is orthogonal such that *UU*^*T*^ = *I*_*n*_. The elements in the diagonal matrix **Λ** are the eigenvalues of ***K***.

By the eigen-decomposition transformation, we have a new random vector 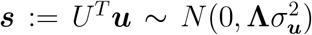, such that **Λ** = *U*^*T*^***K****U*. This transformation immediately results the likelihood in Equation (10) to be

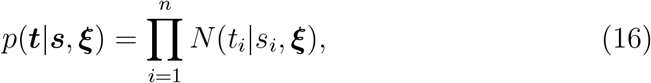

where ***t*** := *U*^*T*^ ***y***. In the way, when updating ***ξ*** in Equation (13) we only need to use a diagonal matrix **Λ** with eigenvalues from ***K*** instead of calling the variance-covariance matrix ***K*** in MCMC or alternative methods.

GEBV:

The genomic breeding values (GEBV) in Equation (1) can be estimated by summing up all the breeding values at each locus *j* from the training population by

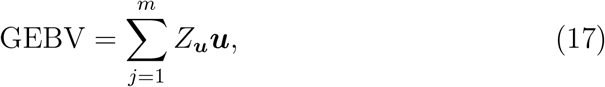

### 3.4. Model scenarios

We examined the value of different explanatory variables, genomic (G), transcriptomic (T), environment (E), their interactions (*G* × *E, G*#*G*) and alternate model frameworks (linear/nonlinear) for prediction of flowering time and height, structuring our analyses across thirteen model scenarios. We applied the traditional GBLUP model as our benchmark as follows:

#### 3.4.1. Models 1-3, additive genomic and non-additive genomic random effects (epistasis and dominance)

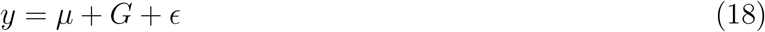

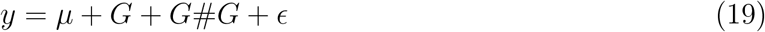

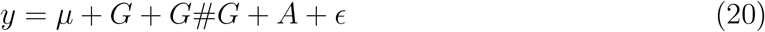

#### 3.4.2. Models 4-5, additive genomic plus genomic and environmental random effects

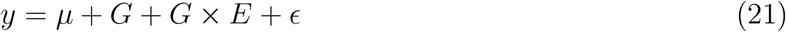

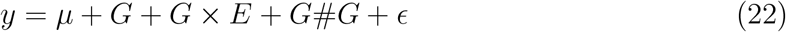

#### 3.4.3. Models 6-7, additive transcriptomic and genomic plus non-additive genomic random effects

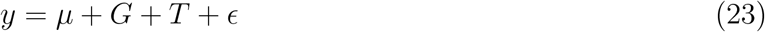

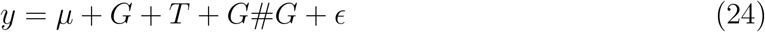

#### 3.4.4. Models 8-9, plus all interaction between omic and environmental random effects

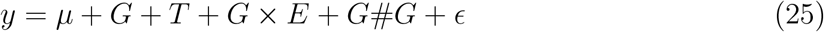

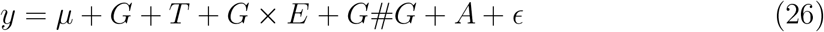

Because of the non-linearity of the Gaussian kernel genomic additive and non-additive random effects are captured together with *G*^*^, which includes G, epstasis (EPI, G#G) and dominant (A), thus we only compare the following

RHKS models with the GBLUP benchmark:

#### 3.4.5. Models 10-11, genomic, transcriptome random effect and their interaction

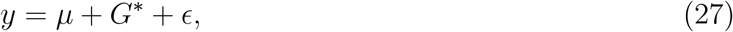

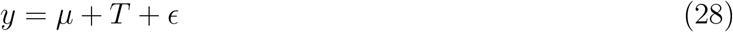

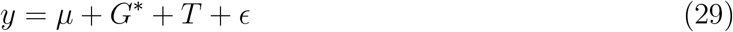

#### 3.4.6. Model 12, plus environmental interactions

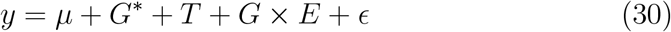

### 3.5. Model assessment

We evaluated model predictive accuracy by 5-fold cross validation (CV, e.g., Allen (1974)) as we do not know the ground truth. This is executed by randomly partitioning 5 equal sized sub samples from the whole population.

In the CV procedure, each sub sample is around 20% of the test population which will be applied to validate the respective model splits trained on the remaining data, around 80% of the population, repeated 5 times. Overall model accuracy was obtained by taking the average of each of the 5 fold model accuracies.

We compute Pearson correlation, *r* by Equation (e.g., Fitch (1984)) between the true trait value ***y*** from the test population and the predictive traits *ŷ* from the predictive model to evaluate model predictive accuracy.

The scheme will

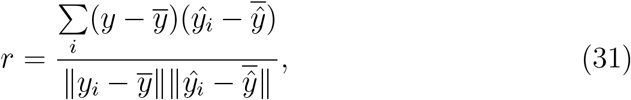

In this work we studied the impacts of genomic additive and non-additive random effects, omics data as well as G × E on the phenotypic traits over two different day lengths for improving GP and GS in for wheat traits. We construct a Bayesian hierarchical model by estimating the parameters of interest in order to compare the predictive models listed in Section 4.2. We chose Gaussian prior for the unknown random effects ***u*** and conjugate priors for ***ξ***, specifically we assign the scaled-inverse *χ*^2^ to all the variance parameters whose hyperparameters were fixed values from R software package, Bayesian generalized linear regression (BGLR, see more details in Pérez and De Los Campos (2014)). The aforementioned models were trained by optimising the unknown parameters using MCMC, which is done by simulating each parameter using the Gibbs sampler from its full conditional posterior distribution formed by the likelihood in Equation (16) after the eigen-decomposition transformation and the selected prior. We also call software package, BGGE, to compute *G* × *E* in Equation (3.2). All our work were implemented by R software.

## 4. Results

In controlled environments, where traits were measured under contrasting long and short daylength regimes, transcriptome abundance (T) outperformed genomic SNPs (G) when they were modelled without environment in the GBLUP regression framework. But the relative improvement afforded by the transcriptome was greater for flowering time than height. This general trend is consistent with the hypothesis that as an intermediate state between the genome, environment and final phenotype, the transcriptome should be a good predictor of trait variation. While the model combining SNP and *G* × *E* effects-marginally outperformed the transcriptome for both traits (Figure 2 and 3). The best performing model for both flowering time and height combined all data types, *G* × *E* and epistasis in the GBLUP and RHKS frameworks, with GBLUP giving slightly better performance overall compared with RKHS across all tests.

**Figure 2.**
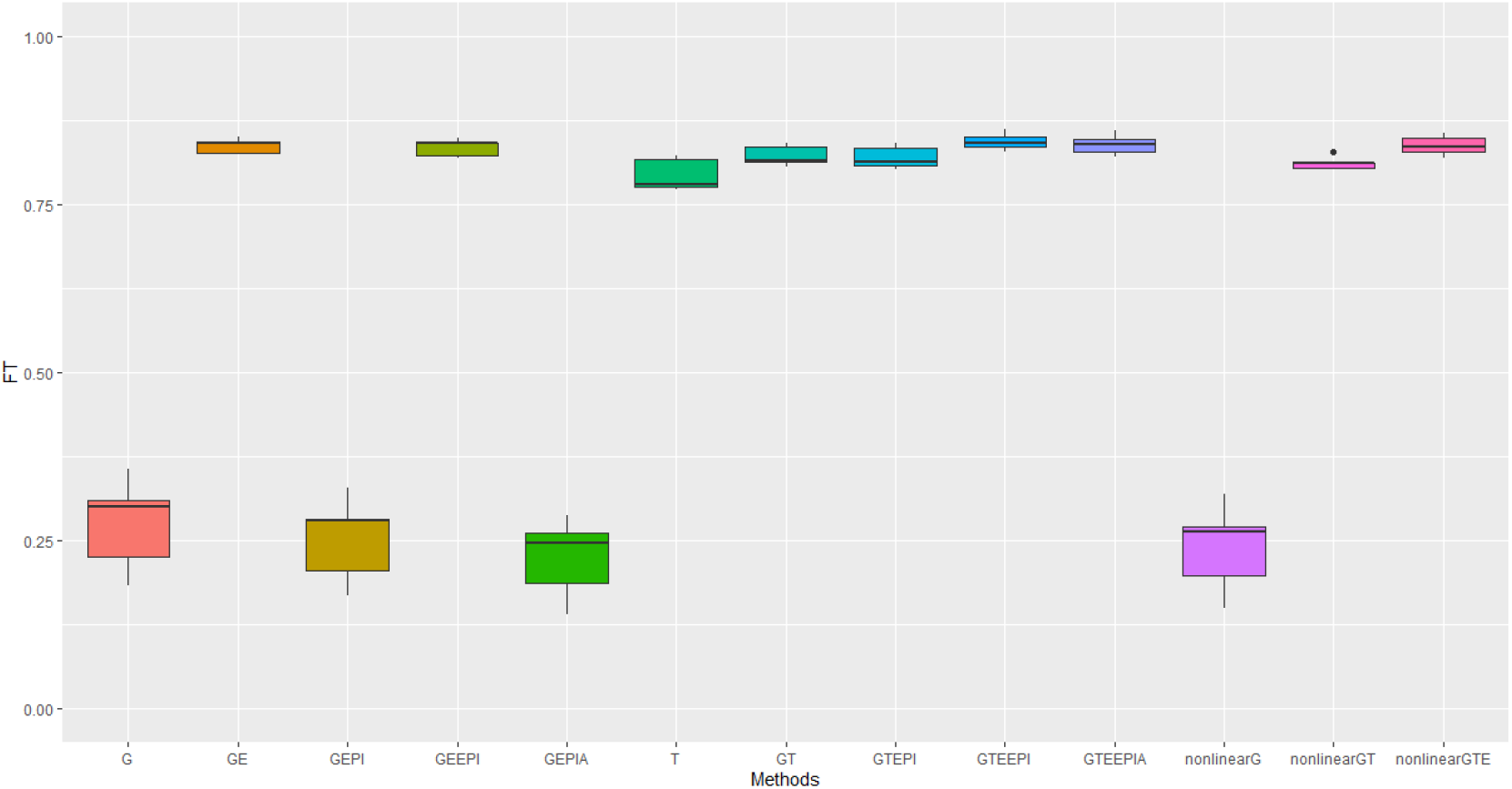
Performance Metrics (Pearson correlation) for genomic predictive accuracy on **FT** across two regression models, GBLUP and RKHS (nonlinear) from controlled environmental data. The error bars represent the model performance among thirteen different model scenarios. The x-axis gives the name of each model in short that is consistent with the described model scenarios in details: **GBLUP**: G, G+GxE (GE), G + G#G (EPI) + A (dominant) (GEPIA), G+GxE+ G#G (GEEPI), G+T (GT), G+T+ G#G (GTEPI), G+T+GxE+ G#G (GTEEPI), G+T+GxE+ G#G +A (GTEEPIA); **RHKS**: G+ G#G + A (nonlinearG), T(T), G+ G#G+ A + T (nonlinearGT), G+ G#G+ A + T + GxE (nonlinearGTE).

**Figure 3.**
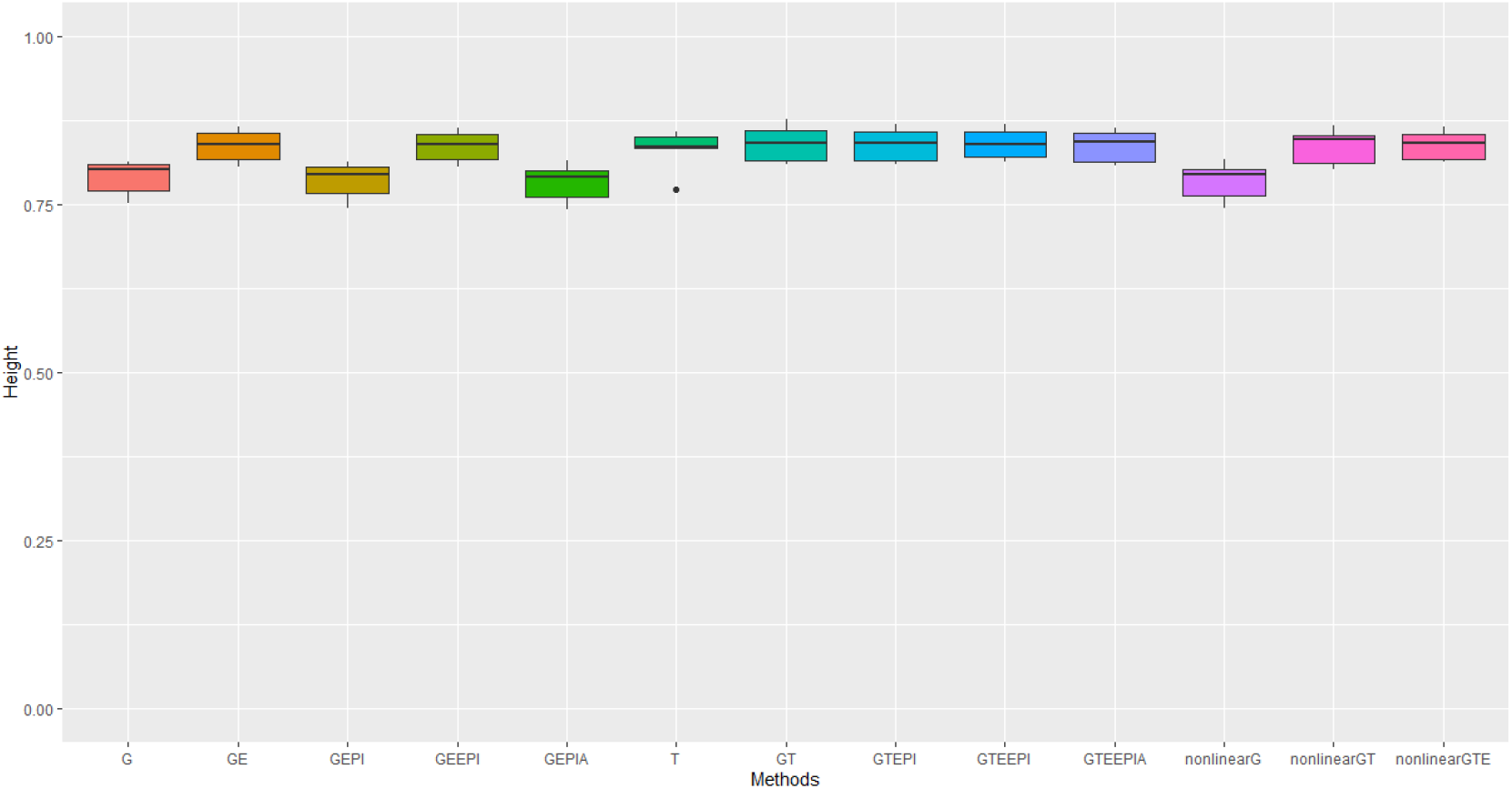
Performance Metrics (Pearson correlation) for genomic predictive accuracy on **height** across two regression models, GBLUP and RKHS (nonlinear) from controlled environmental data. The error bars represent the model performance among thirteen different model scenarios. The x-axis gives the name of each model in short that is consistent with the described model scenarios in details: **GBLUP**: G, G+GxE (GE), G + G#G (EPI) + A (dominant) (GEPIA), G+GxE+ G#G (GEEPI), G+T (GT), G+T+ G#G (GTEPI), G+T+GxE+ G#G (GTEEPI), G+T+GxE+ G#G +A (GTEEPIA); **RHKS**: G+ G#G + A (nonlinearG), T(T), G+ G#G+ A + T (nonlinearGT), G+ G#G+ A + T + GxE (nonlinearGTE).

**Figure 4.**
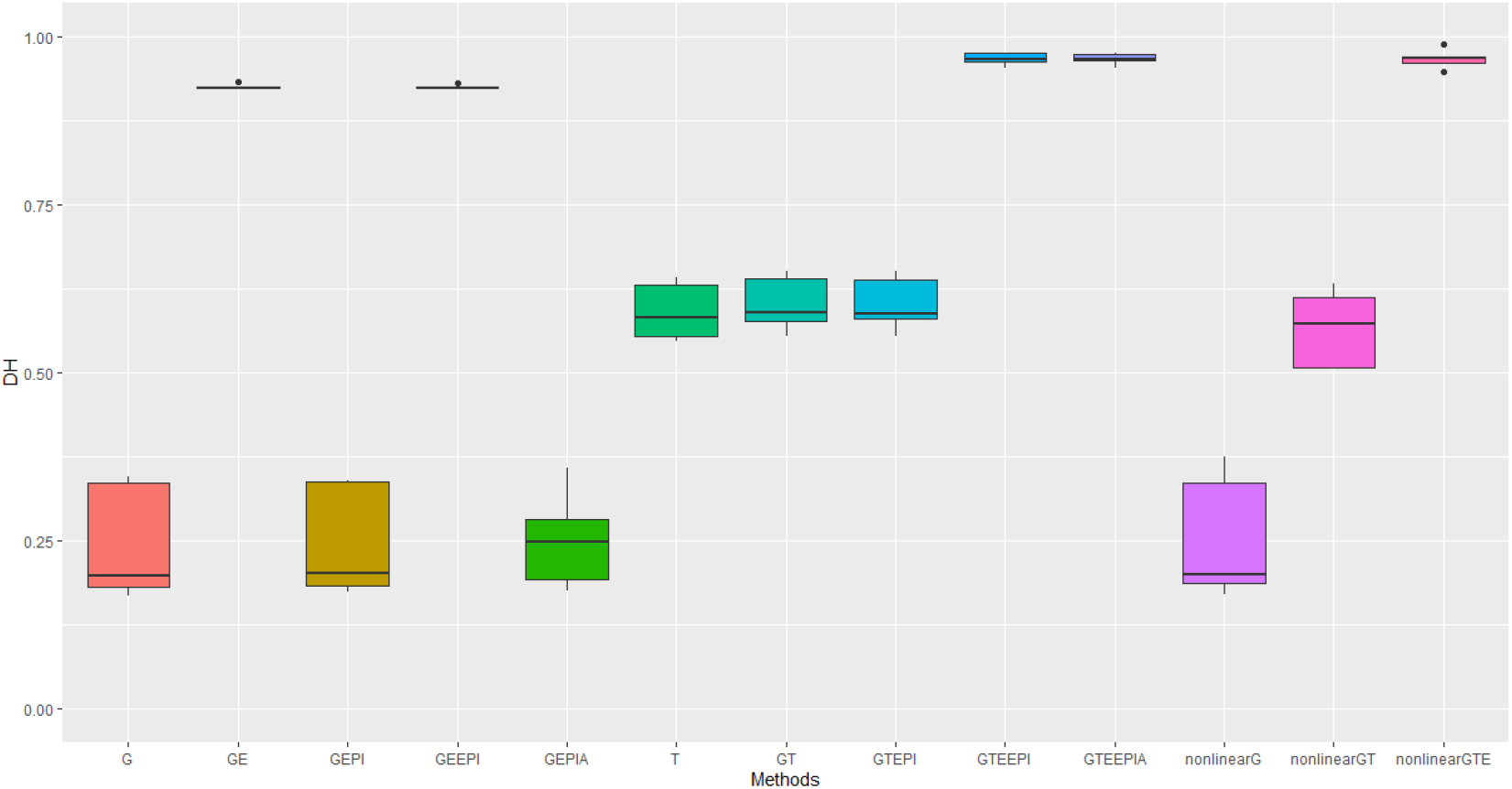
Performance Metrics (Pearson correlation) for genomic predictive accuracy on **DH** across two regression models, GBLUP and RKHS (nonlinear) from field experiment data. The error bars represent the model performance among thirteen different model scenarios. The x-axis gives the name of each model in short that is consistent with the described model scenarios in details: **GBLUP**: G, G+GxE (GE), G + G#G (EPI) + A (dominant) (GEPIA), G+GxE+ G#G (GEEPI), G+T (GT), G+T+ G#G (GTEPI), G+T+GxE+ G#G (GTEEPI), G+T+GxE+ G#G +A (GTEEPIA); **RHKS**: G+ G#G + A (nonlinearG), T(T), G+ G#G+ A + T (nonlinearGT), G+ G#G+ A + T + GxE (nonlinearGTE).

**Figure 5.**
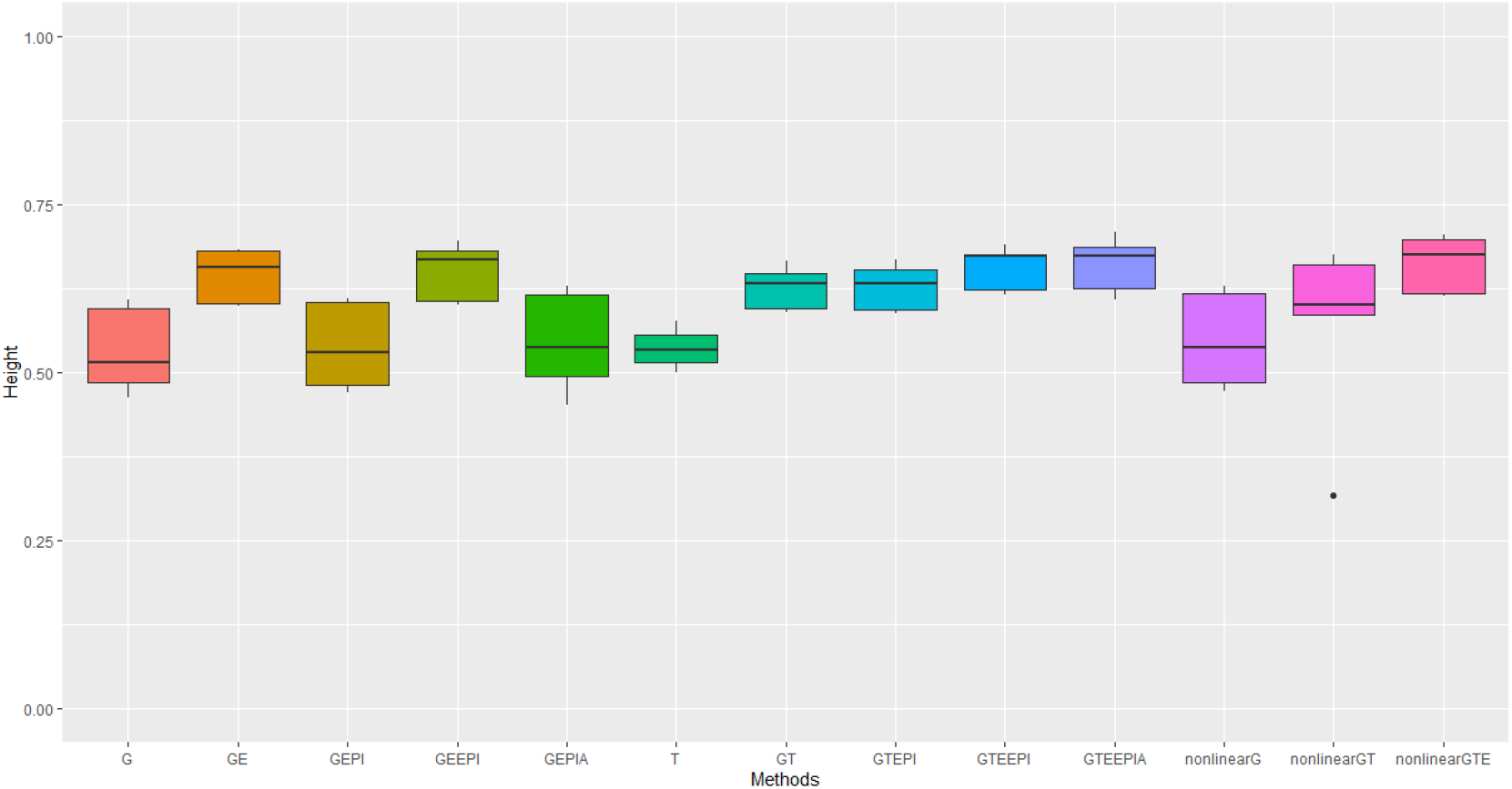
Performance Metrics (Pearson correlation) for genomic predictive accuracy on **height** across two regression models, GBLUP and RKHS (nonlinear) from field experiment data. The error bars represent the model performance among thirteen different model scenarios. The x-axis gives the name of each model in short that is consistent with the described model scenarios in details: **GBLUP**: G, G+GxE (GE), G + G#G (EPI) + A (dominant) (GEPIA), G+GxE+ G#G (GEEPI), G+T (GT), G+T+ G#G (GTEPI), G+T+GxE+ G#G (GTEEPI), G+T+GxE+ G#G +A (GTEEPIA); **RHKS**: G+ G#G + A (nonlinearG), T(T), G+ G#G+ A + T (nonlinearGT), G+ G#G+ A + T + GxE (nonlinearGTE).

In terms of non-additive interaction effects, the inclusion of *G* × *E* had the most positive impact on prediction accuracy, in particular for flowering time. Prediction of height was less dependent on interactions with environment and hence the G and G+*G* × *E* models performed better relative to anthesis. Explicitly fitting epistatic (EPI) and dominance (A) interactions based on SNP covariance slightly reduced model accuracy under the GBLUP for both agronomic traits. While we cannot test this trend precisely with RHKS because G and *G*#*G* cannot be disentangled, the RHKS model scenarios including non-additivity outperformed their GBLUP equivalents.

Predictive accuracy for height and days to heading measured in four field experiments over 2 years suggest similar outcomes for the tested model scenarios as observed in controlled environments, with the following exceptions. Under field conditions predictive accuracies were lower for height across all model scenarios. Whereas predictions for flowering for the best performing scenarios were higher than in controlled environments. In particular the improvement gained from including *G* × *E* into predictions for flowering was substantial and improved on that observed in controlled environments. Of course, any direct comparisons need to be treated carefully here. The environment contrasts in controlled (daylength) and field (rainfall and temperature) experiments are based on different underlying variables which potentially could explain some differences including the magnitude of improvement from including *G* × *E* under both the GBLUP and RHKS regression.

The transcriptome on its own again performs better than G for predicting flowering but the relative improvement is not as large as seen in controlled environment experiments (Table 2). Whereas for height there was no discernible advantage to including the transcriptome over SNP data alone, which underperformed relative to the model scenarios including G and *G* × *E* under the GBLUP framework. Inclusion of non-additive effects (*G* × *E*, epistasis and dominance) under the GBLUP framework all improved model accuracy, counter to daylength experiments. The improvement to prediction of both traits, but flowering, through inclusion of *G* × *E* remained substantial relative to epistasis and dominance effects. As with the controlled environments RHKS outperformed the GBLUP equivalents. Finally, in contrast to controlled environments the RHKS framework, for the best performing models combining all data types, outperformed or performed equally well to the equivalent GBLUP model.

In summary, the most predictive models were those which combined all data types. In both sets of experiments the transcriptome is predictive of both traits, but much more so in controlled environments. This is somewhat expected because gene expression better captures how genes contribute to trait development under different environments, see e.g., Michel et al. (2021). Inclusion of the transcriptome potentially helps capture key genes and pathways involved in trait expression, which can refine predictive models. The nonlinear regression model did not show many advantages than the traditional linear mixed models from these two data sets, especially for the controlled environmental data. It outperformed or was similar to GBLUP in field conditions, possibly reflecting that in controlled environments, genotype and environment effects and interactionss are less complex and more easily delineated, allowing for the isolation of these variables from more complex environmental interactions, with implications for suitability of different model frameworks. In both data sets, the site/treatment covariates simplify the environmental variation, which can reduce the impact of complex non-additive genomic interactions that are more pronounced in variable natural conditions, see e.g., Teressa et al. (2021); Becker and Leon (1988).

## 5. Discussion

Crop breeding is at a pivotal point. The combined factors of expanding population growth, limited availability of arable land for agriculture and changing environment are demanding improved avenues for profitable, sustainable crop production worldwide. The industry needs solutions that can provide more accurate, high-throughput decision making to fast-track development of improved crop cultivars, targeting staple crop genetics, environments and management practices. Modern statistical breeding technologies including new and improved avenues for integrating multiple objectives into genetic evaluations are fundamental for decision making.

We examined the potential of the transcriptome as an alternative predictor in genomic selection (GS) for wheat developmental traits, flowering time and height. The potential for the transcriptome to add value as a predictor in GS has steadily gained attention in recent years (Xu et al., 2017). Recent studies have explored the transcriptome’s predictive value in GS for crops such as maize (Zheng et al., 2017; Schrag et al., 2018; Azodi et al., 2019) and rice (Wang et al., 2019), but its application in wheat, one of the world’s major food crops, remains under explored. Furthermore, most studies focus on controlled conditions, whereas applying omics data in real-world settings is crucial for assessing its utility in practical breeding programs. Consistent with previous findings, we found the transcriptome to be an effective phenotype predictor within both GBLUP and RHKS regression models, particularly outperforming genomic SNPs for anthesis and heading date, and for height in controlled conditions, when environment was excluded from the model.

The underperformance of genomic SNP models in predicting flowering time likely stems from the trait’s strong dependence on genotype-environment interactions, such as daylength and temperature (Susila et al., 2018; Ausin et al., 2005). This effect may have been more pronounced under non-vernalising conditions in controlled environments, due to interactions between the VRN1 locus, which can delay flowering in vernalisation sensitive lines, and photoperiod sensitive genes (PPD1) and the FT locus under different day length conditions (Hyles et al., 2020). Excluding environmental factors as in Model scenarios 1-3, 10, which only considered genetic factor in isolation, led to poor cross-validation accuracy in genomic prediction. Indeed, when *G* × *E* was included, a much greater proportion of the variation in flowering was explained by the model (see Table 1 and 2). In addition, the transcriptome’s improved performance suggests it captures these genotype-environment interactions, making it a more reliable predictor of flowering time. This aligns with the hypothesis that the transcriptome, as an intermediary between the genome, environment, and phenotype, effectively captures these effects and should be a reliable predictor of trait variation (Te Pas et al., 2017).

**Table 1:** Model predictive accuracy and relative uncertainties.

| Controlled environment |  |  |
| --- | --- | --- |
| Model | Height(cm)<br>(std dev) | Athesis(DAS)<br>(std dev) |
| (1) G | 0.789 (0.027) | 0.275 (0.070) |
| (2) G+G#G (EPI) | 0.785 (0.029) | 0.268 (0.058) |
| (3) G+G#G +A (dominant) | 0.782 (0.030) | 0.253 (0.064) |
| (4) G+G×E | 0.837 (0.025) | 0.837 (0.011) |
| (5) G+G×E+ G#G | 0.836 (0.024) | 0.834 (0.013) |
| (6) G+T | **0.840 (0.030)** | 0.822 (0.016) |
| (7) G+T+G#G | 0.838 (0.026) | 0.819 (0.017) |
| (8) G+T+G×E+G#G | **0.840 (0.023)** | **0.843 (0.013)** |
| (9) G+T+G×E+G#G+A | 0.836 (0.025) | 0.839 (0.015) |
| (10) RHKS: G+G#G+A | 0.784 (0.030) | 0.240 (0.067) |
| (11) RHKS: T | 0.836 (0.028) | 0.811 (0.011) |
| (12) RHKS: G+G#G+A+T | 0.836 (0.028) | 0.811 (0.011) |
| (13) RHKS: G+G#G+A+T+G×E | 0.838 (0.023) | 0.837 (0.015) |

Pearson correlation by Equation (31) was used to indicate predictive accuracy of model performance from the controlled experiment data. The mean and standard deviation (shorten by "std dev" in the bracket) of Pearson correlations were calculated from 5-fold cross validation between training and test populations based on thirteen different model scenarios that described in Section 3.4.

**Table 2:** Model predictive accuracy and relative uncertainties.

| Field experiments |  |  |
| --- | --- | --- |
| Model | Height(cm)<br>(std dev) | Days to Heading (std dev) |
| (1) G | 0.533 (0.065) | 0.245 (0.087) |
| (2) G+G#G (EPI) | 0.539 (0.066) | 0.247 (0.084) |
| (3) G+G#G+A | 0.545 (0.076) | 0.251 (0.073) |
| (4) G+G×E | 0.644 (0.041) | 0.925 (0.005) |
| (5) G+G×E+G#G | 0.650 (0.044) | 0.924 (0.004) |
| (6) G+T | 0.625 (0.033) | 0.602 (0.042) |
| (7) G+T+G#G | 0.627 (0.036) | 0.603 (0.041) |
| (8) G+T+G×E+G#G | 0.656 (0.033) | **0.967 (0.009)** |
| (9) G+T+G×E+G#G+A | **0.660 (0.042)** | **0.966 (0.009)** |
| (10) RHKS: G+G#G+A | 0.548 (0.072) | 0.253 (0.095) |
| (11) RHKS: T | 0.536 (0.030) | 0.591 (0.044) |
| (12) RHKS: G+G#G+A+T | 0.558 (0.134) | 0.566 (0.058) |
| (13) RHKS: G+G#G+A +T+G×E | **0.662 (0.044)** | **0.966 (0.015)** |

Pearson correlation by Equation (31) was used to indicate predictive accuracy of model performance from field experiment data. The mean and standard deviation (shortened by "std dev" in the bracket) of Pearson correlations were calculated from 5-fold cross validation between training and test populations based on thirteen different model scenarios that described in this work, Section 3.4.

In the case of height we see a different story. While inclusion of *G* × *E* does improve model predictive accuracy in controlled conditions and the field, the relative gain is reduced compared to flowering time. Suggesting that simpler genetics underpinning height, predominantly major genes from the Rht family (Zheng et al., 2017; Achard et al., 2009). In the field there was no advantage to using the transcriptome over genome SNPs. This could be explained by the fact that these key height controlling genes are not expressed until later in development (Borrill et al., 2022), hence transcriptomes collected at the earlier stage in this study would be unlikely to capture GxE effects at these loci.

Although the transcriptome offers predictive benefits for flowering time, particularly in controlled environments, its effectiveness in field conditions is reduced. This likely reflects the incompleteness with which a transcriptome taken at a single time point early in development can capture *G* × *E* effects experienced throughout development to maturity in the field. The controlled environment experiments would not suffer this limitation, as environmental conditions were maintained throughout development, preserving the relationship between the regulatory signal captured in the early-stage transcriptome and the trait. This highlights a challenge in using highly plastic omic data, like the transcriptome, for GS in variable environments, where multiple tissues and time points might be needed to capture relevant interactions, complicating its application in commercial breeding programs. Nonetheless, it is notable that the early-stage transcriptome offers some predictive power for flowering time in field conditions.

Our findings indicate that while the inclusion of the transcriptome or direct modelling of genotype-environment interactions is essential for reliable predictions, especially for flowering time, other non-additive factors such as epistasis and dominance made only minor contributions. Consistent with expectations (Gianola and Van Kaam, 2008), RHKS was slightly better at capturing complex genomic random effects compared to the GBLUP model (e.g. Model scenarios 13 and 9; 10 and 3), although the advantage was marginal.

The best-performing models for both traits combined all data types, genotype-environment interactions, and epistasis under the GBLUP framework in controlled conditions and the RHKS framework in the field. The RHKS model’s marginal advantage in field conditions may reflect greater complexity of environmental variables that the non-linear Gaussian kernel can better capture, see e.g., Cuevas et al. (2017). However, in controlled environments where environments are simplified and therefore the *G* × *E* interactions likely to be less complex than in the field, the GBLUP model is promising. Despite the transcriptome’s predictive advantages, the high cost and complexity of incorporating it into breeding programs currently limit its practicality. Additionally, the need to structure sampling around developmental and environmental cues for effective trait prediction adds another layer of complexity, which affects the utility of omics data for genomic selection (GS) in commercial breeding programs. For the time being, the strength of population-scale transcriptomics lies in enhancing our biological understanding of complex genomic interactions, which can then be integrated into breeding selection models in alternative ways (Khalilisamani et al., 2024).

For both flowering time and height, incorporating genotype-environment interactions into genomic SNP models provided comparable prediction accuracy in both controlled and field conditions, suggesting a more practical approach for improving GS when environmental covariates are well characterised. Recent studies (Varona et al., 2018; Morais Junior et al., 2017) have supported this approach, showing improved prediction accuracy with inclusion of genotype-environment modelling, which could enhance genetic evaluations for breeding programs across diverse environments.

In conclusion, in both controlled and field experiments, transcript data performs well relative to SNP or environment in isolation for predicting plant height and flowering time due to its ability to capture both genetic and environmental signals and their interactions. By integrating the transcriptome with genetic SNPs and *G* × *E* interactions, models provide a wholistic and highly accurate solution to predict these complex traits. However, the present cost of this combined intervention remains infeasible compared to predictive gains. In light of this we point to the inclusion of *G* × *E* and epistasis in the GS framework as a potentially more reliable substitute if environments can be well characterised.

## 6. Acknowledgements

This research was led by the CSIRO, funded through the Agriculture and Food Strategic Investment and the Machine Learning and Artificial Intelligence Future Science Platform. Communication relating to this work should be directed to CSIRO.

We thank Dr Hyles Jessica for providing insightful comments on the manuscript, which improved the biological interpretation. We thank Dr Ben Trevaskis for sharing his knowledge and experience to help this work.

We generously thank the many volunteers at CSIRO and NSW-DPI who assisted with field campaigns to collect transcriptome samples in GES and WAGGA, in particular, Emmet Leynne, Aswin Singaram Natarajan, Alex Boyer, Jessica Hyles, Bjorg Sherman, David Deery, Marck Cmiel, Saul Newman, Trijntje Hughes, Todd Collins, Hayden Petty, Cameron Copeland, Dean Mccallum, Javier Atayde, but many more.

## 7. Data Assess

The field data that analysed in this work will be release together with this work once it was accepted.

